# Transcriptome features of trained immunity in *Drosophila*

**DOI:** 10.1101/2021.12.22.473822

**Authors:** Naoyuki Fuse, Chisaki Okamori, Chang Tang, Kikuko Hirai, Ryoma Okaji, Shoichiro Kurata

## Abstract

Immune memory is an ability of organisms to potentiate immune responses at secondary infection. Current studies have revealed that innate immunity, as well as adaptive immunity, exhibits the memory character called “trained immunity”. Although it is suggested that epigenetic reprogramming plays important roles in trained immunity, its underlying mechanism is not fully understood, especially on the individual level.

Here we established experimental systems for detecting trained immunity in *Drosophila melanogaster*. Namely, training infection with low-pathogenic bacteria enhanced the survival rate of the flies at subsequent challenge infection with high-pathogenic bacteria. We found that among low-pathogenic bacteria, *Micrococcus luteus* (Ml) and *Salmonella typhimurium* (St) mediated apparent training effects in fly, but seemed to act through different ways. Ml left training effects even after its removal from flies, while living St persisted inside flies for a long time. Our RNA-Seq analysis revealed that Ml-training enhanced the expression of immune-related genes during the challenge infection, but did not do so without challenge infection. In contrast, St-training maintained high expression of the immune-related genes with or without challenge. These results suggest that training effects with Ml and St were due to memory and persistence of immune responses, respectively. Furthermore, we searched the factor involved in Ml-training and identified a candidate, Ada2b, which is a component of the histone modification complex. We found that the *Ada2b* RNAi and mutant flies showed dampened enhancement of survival rates after Ml-training. These results suggest that Ada2b is involved in the *Drosophila* trained immunity.

## Introduction

Immunity is necessary for organisms to fight against pathogens. The immune machinery generally consists of two systems: innate immunity and adaptive immunity. Innate immunity is the primitive immune system evolutionarily conserved among multicellular organisms and functions at the first line of the defense. Since this system recognizes the molecular patterns of pathogens, its reaction is not specific to a pathogen but works in heterologous manners. In contrast, adaptive immunity recognizes pathogens in highly specific manners and was evolutionarily developed in the lineage of vertebrates. Moreover, adaptive immunity exhibits the character of memory that potentiates immune responses at secondary infection. Although it has been thought for a long time that innate immunity does not possess memory character, recent studies have provided evidence against this concept [1,2].

Bacillus Calmette Guérin (BCG) vaccine has been widely used for protecting infants from *Mycobacterium tuberculosis*. During its long history, it has been frequently observed that BCG vaccination promotes the protection against not only tuberculosis but also a wide variety of pathogens in heterologous manners. Recent studies demonstrated that the innate immune system is responsible for heterologous effects of the BCG vaccine [3,4]. Moreover, the SCID mouse deficient for the adaptive immune system preserves the capacity to enhance immune responses at subsequent infection [5], suggesting memory functions of innate immune system. Thus, the heterologous reinforcement of immunity through repeated infections has been observed in many other organisms and systems [6– 8]. These phenomena are called “innate immune memory” and also “trained immunity”[1,2]. As seen in the case of BCG vaccine, “trained immunity” has attracted notice in the clinical field, and many researchers are addressing the molecular mechanisms underlying it.

The innate immune cells of mammals (e.g., macrophages and monocytes) respond to the pathogen-associated molecular patterns (PAMPs) and change their gene expression in vitro, as well as in vivo. Previous studies showed that the responses of gene expression upon primary stimulus were enhanced or suppressed upon secondary stimulus, suggesting “potentiation” or “tolerance” of immune responses, respectively [9–13]. Anti-microbial peptide (AMP) genes, for example, showed potentiation of gene expression during secondary stimulus, while pro-inflammatory genes (e.g., interleukins) showed tolerance of gene expression. It is thought that these responses of gene expression represent the trained immunity at the molecular level. Furthermore, it is shown that epigenetic regulation of gene expression plays important roles in trained immunity. Histone modifications, such as methylation of histone H3 Lys4 (H3K4) and acetylation of histone H3 Lys14 (H3K14), are dynamically changed by immune stimuli, and modify the gene expression in response to secondary stimuli [10–13]. Epigenetic reprogramming is observed not only in short-lived peripheral myeloid cells (e.g., macrophages and monocytes) but also in long-lived stem cells (e.g., hematopoietic stem and progenitor cells: HSPCs) [14,15], consistent with long-lasting effects of the trained immunity in mammalian individuals. Moreover, metabolic shifts of cholesterol synthesis and glycolysis in innate immune cells are observed under immune training, and contribute to the potentiation of immune responses [16,17]. Thus, numerous data have been accumulated in the field of “trained immunity”, but their mechanisms are not fully understood yet. For example, it is not known how immune information is converted into epigenetic information during trained immunity. It is also unclear how trained immunity is organized on the individual level.

We study “trained immunity” using *Drosophila*. This insect is amenable to various genetic tools for dissecting the molecular mechanisms on the individual level. Insects do not possess adaptive immunity, and protect themselves from pathogens only via the innate immune system. Therefore, analysis of innate immunity in insects is relatively simple and has untangled its molecular networks this decades [18,19]. Molecules involved in innate immunity are well conserved between insects and mammals. For example, the Toll and Imd pathways are two major signaling pathways of *Drosophila* immunity, and they correspond to Toll-like receptor (TLR) and tumor necrosis factor receptor (TNF-R) signaling pathways in mammals, respectively. These signaling pathways in *Drosophila* display some broad specificity against pathogens. The Toll pathway is activated mainly by gram-positive bacteria and fungi, while the Imd pathway is activated mainly by gram-negative bacteria. However, these responses are not exclusive, but rather are overlapping to some extent. Ultimately, these signaling pathways regulate the expression of immune-related genes, such as AMPs, through NF-κ B transcription factors.

In response to infection, AMPs are produced and secreted mainly from the fly fat body (corresponding to the mammalian liver) into hemolymph (body fluid), and those attach directly to pathogens [18,19]. In addition to this humoral response, the cellular response of immunity is mediated by fly hemocytes (corresponding to mammalian macrophages). Hemocytes incorporate pathogens into cellular vesicles through phagocytosis to kill them. One kind of hemocytes also induces coagulation and melanization of hemolymph to prevent the proliferation of pathogens. Humoral and cellular immunity are organized locally and systemically. Moreover, non-immune cells, such as gut, muscle, neuronal, and reproductive organ cells also respond to pathogen infection [20]. These responses are coordinated on the individual level and influence physiology, metabolism, behavior, and homeostasis of fly individuals.

We still do not know how trained immunity is evolutionarily conserved among organisms. In *Drosophila*, phenomena related to trained immunity have been observed in several previous experiments [21–23]. In those experiments, primary infections enhanced the survival rates of flies after secondary infections. However, these training effects on survival rate could be attributed to immune persistence rather than immune memory [9]. In the case of immune persistence, primary immune activation is maintained at the time of secondary infection. In the case of immune memory, primary immune activation has ceased at the time of secondary infection but reinforces immune responses against secondary infection. It remains obscure which mode acts in the “trained immunity” in fly.

Here, we established experimental systems of trained immunity in *Drosophila*. We performed RNA-Seq analysis and observed memory and persistence of immune responses at the molecular level. We also identified a chromatin regulator potentially involved in the trained immunity in fly.

## Materials and Methods

### Flies

The *Drosophila melanogaster* lines used in this study were Oregon-R (Bloomington *Drosophila* Stock Center (BDSC) #6362), da-Gal4 (BDSC#55851), UAS-GFP RNAi (BDSC#9330), UAS-*Ada2b* RNAi (National Institute of Genetics, Japan (NIG) #9638R-3), *white*[1118] mutant (BDSC#3605), *Ada2b* [d272] mutant (a gift from Dr. N. Zsindely, University of Szeged) [24, 25] lines. Flies were reared on standard corn meal medium at 25 ºC.

### Bacteria

We used the following bacteria in this study: *Micrococcus luteus* (Ml: IFO:13276), *Salmonella typhimurium* (St: SL1344), *Staphylococcus saprophyticus* (Ss: GTC:0205), *Erwinia carotovora carotovora 15-GFP* (Ec) [26], *Staphylococcus aureus* (Sa: ATCC10801), *Pseudomonas aeruginosa* (Pa: ATCC15692), Sa-GFP (RN4220 / pTetON-GFPopt, spectinomycin-resistant strain) [27], Pa-Kan (PA01 / pBBR1MCS2, kanamycin-resistant strain) [28], and Sa-Lux (ATCC 12600 / Tn4001-luxABCDE-kan, kanamycin-resistant strain) [29]. Ml, St, Ec, Sa, and Pa bacteria were cultured in LB medium (Nacalai Tesque), and Ss was cultured in NB medium (Becton Dickinson). Sa-GFP and Pa-Kan were cultured in medium with antibiotics: spectinomycin (100 µg/ml) and kanamycin (200 µg/ml), respectively. Sa-Lux was cultured in TSB medium (Becton Dickinson). Ml was cultured at 30 ºC, and other bacteria were cultured at 37 ºC.

Bacteria used for training were cultured overnight (∼16 hours) and were pelleted by centrifugation (5000 rpm, 10 minutes). Bacteria pellets were resuspended in saline (Otsuka Pharmaceutical), and their concentrations were adjusted to the concentrations indicated in the text (usually OD = 1, unless otherwise noted). Bacteria used for challenge were re-cultured for 3 hours from a 1: 100 dilution of the overnight culture. Challenge bacteria were also precipitated and diluted to the concentrations indicated in the text (Pa and Pa-GFP: OD = 1E-5, Sa: OD = 0.2, Sa-GFP and Sa-Lux: OD=0.1, unless otherwise noted) in saline.

### Survival assay

Infection experiments were performed as described previously [30]. Briefly, adult male flies aged from 4 to 7 days were used for the experiments. Bacteria solution or saline (control) was drawn into a glass needle (Drummond, 3 1/2 inch capillary), and 70 nl of the solution was injected into body cavity of the fly thorax by using a micromanipulator (Drummond, Nanoject II). Challenge injection was usually performed at 6 days after training injection, unless otherwise noted. The flies that died within 3 hours after challenge were ignored for the survival assay. From the next day, dead flies were counted every 2 hours for Pa challenge and every day for Sa challenge. Statistical comparisons of survival curves were performed by Log-rank test and post-hoc Tukey HSD test using the R program.

### Bacteria load assay

For measuring the bacteria load, the flies at the indicated times after injection were collected and were washed briefly with 70 % ethanol. Then one fly was placed into a tube and was homogenized using a pestle in 100 µl of LB medium. After making serial dilutions of fly extracts, 10 µl of diluted extracts were spotted on LB agar plates, and the plates were incubated at 30 ºC (Ml) or 37 ºC (St, Sa, Pa) overnight. Bacteria load of the original fly extract (colony formation units per fly: cfu / fly) was calculated according to colony numbers and dilutions. In this protocol, we did not observe any colonies from control flies without training or challenge. For measuring the bacteria load of challenge, antibiotic-resistant strains of challenge bacteria (Sa-GFP and Pa-Kan) were used, and fly extracts were spotted on LB agar plates containing antibiotics as described above. Statistical comparisons of bacteria loads were performed by Kruskal-Wallis ANOVA and post-hoc Wilcoxon rank sum test using the R program.

### Sample preparation and RNA-Seq

We performed training and challenge infections of Oregon-R adult male flies as described above. The flies at 4 hours after challenge were collected in tubes, were quickly frozen in liquid nitrogen, and were stored at -80 ºC until RNA extraction. As one sample for RNA-Seq, 10 flies were homogenized in TRIzol reagent (Thermo Fisher Scientific), and total RNAs were extracted according to the manufacturer’s standard procedure. We evaluated the yield and purity of RNAs using Nano-Drop (Thermo Fisher Scientific), Qubit (Invitrogen), and Bioanalyzer (Agilent). The libraries were prepared using a Strand-specific RNA Library Prep kit (Agilent), and the sequencing (36 bases, single-end) was performed using Illumina HiSeq 2000. Raw data of sequences were deposited in DDBJ, DRA (accession number DRA008187).

### Data analysis for RNA-Seq

Transcriptome analyses were performed using the Linux or the Macintosh operating system. Adaptor sequences were removed from read sequences using Cutadapt [31]. Cleaned read sequences were mapped on the *Drosophila melanogaster* reference genome (ver. 6.04) using the Hisat2 program [32] with default parameters. Gene annotation data (ver. BDGP6.79) were utilized to attribute reads to genes. The number of reads was counted for each gene using Htseq [33], and was normalized using edgeR [33]. Multidimensional scaling (MDS) analysis was performed using edgeR and R programs. Differentially expressed genes (DEGs) were identified using a statistical criterion (generalized linear model likelihood ratio test, false discovery rate (FDR)-adjusted p-value < 0.05). Clustering analysis of DEGs was performed using the R program. The web-based databases DAVID [34], FlyMine [35] and FlyBase [36] were used for analyses of Gene Ontology (GOTERM_BP_DIRECT category), publication enrichment, and gene function.

### qPCR analysis

We performed quantitative PCR (qPCR) analysis for the *Ada2b* and *RpL32* genes.

We extracted total RNA from three flies for each sample and analyzed three samples as biological replicates for each experimental condition. cDNAs were synthesized from total RNAs using ReverTra Ace (Toyobo) according to the manufacturer’s protocol. The qPCR reactions were performed in triplicate for each cDNA sample using the Thunderbird Next SYBR qPCR Mix kit (Toyobo) on Light Cycler 96 (Roche). The primers used were the followings: 5’-ATATACACCCGCCGACTCAG-3’ and 5’-GATCCCAATAGCCGCTCATA-3’ for the *Ada2b* gene, and 5’-AGATCGTGAAGAAGCGCACCAAG-3’ and 5’-CACCAGGAACTTCTTGAATCCGG-3’ for the *RpL32* gene. The relative gene expression was quantified using the R program, and data were statistically analyzed by ANOVA and post-hoc Tukey HSD test on the R program.

## Results

### Experimental system for detecting trained immunity

We initially attempted to establish an experimental system to detect trained immunity in *Drosophila*. Considering the effectiveness of live vaccines, we examined the training effects of live bacteria. Namely, low-pathogenic bacteria were injected into wild-type flies (Oregon-R strain) as training, and at day 6 after training, high-pathogenic bacteria were injected into some flies as challenge. Subsequently we measured the survival rates of the flies (Fig. 1A). As controls, saline was injected into flies for training and/or challenge. We tested various combinations of low-pathogenic bacteria (*Micrococcus luteus* (Ml), *Salmonella typhimurium* (St), *Staphylococcus saprophyticus* (Ss), or *Erwinia carotovora carotovora 15* (Ec)) for training and high-pathogenic bacteria (*Staphylococcus aureus* (Sa) or *Pseudomonas aeruginosa* (Pa)) for challenge. Thereby, we observed increased survivals of the trained, challenged flies with several of these combinations compared to control untrained, challenged flies. For example, challenge injection of Sa gradually killed most of the flies within a week, but the advance training with Ml significantly increased the survival of the flies after challenge with Sa (Fig. 1B). Furthermore, Pa was highly pathogenic and killed all flies within a day. Training with St had a drastic effect so that most of the trained flies survived for days after challenge with Pa (Fig. 1C).

**Fig. 1.**
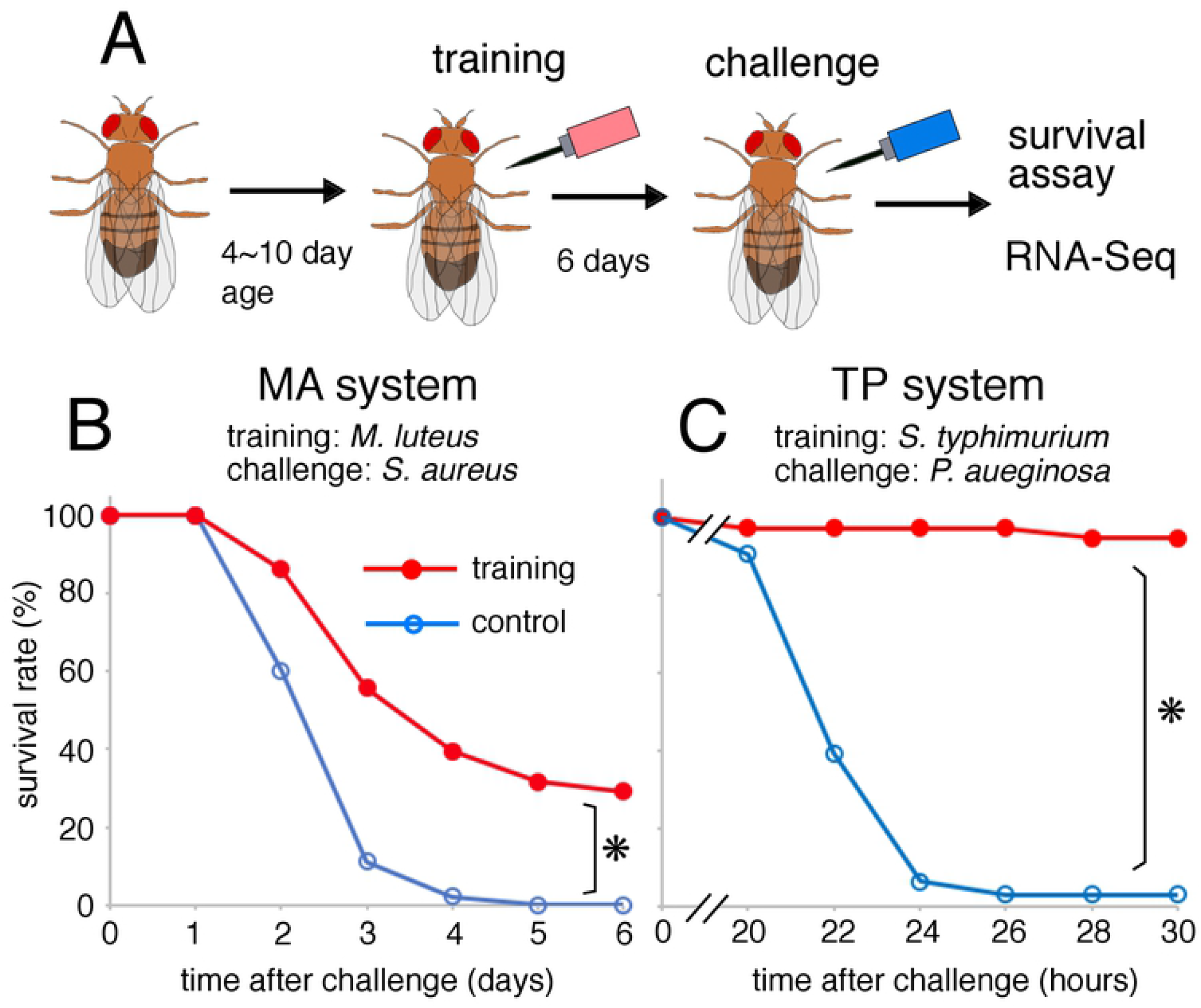
Experimental system of *Drosophila* trained immunity (A) Schematic drawing of systemic infection experiment. Low-pathogenic bacteria were injected into body cavity of flies, and 6 days later, high-pathogenic bacteria were injected. After that, fly survival was monitored. (B) Survival curve of MA system: Ml-training and Sa-challenge. (C) Survival curve of TP system: St-training and Pa-challenge. Red and blue lines represent the flies with and without training, respectively. Asterisks indicate statistically significant (p-value < 0.05) in Log-rank test. Numbers of flies used in these experiments are (B) 40, 57, and (C) 42, 37 (with or without training, respectively).

The training effect seemed not to be specific, but rather showed cross reactivity (S1 Fig). Ml-training significantly increased survival of the flies after Pa-challenge as well as Sa-challenge. Moreover, St showed training effects against Sa as well as against Pa. However, some combinations of bacteria displayed some specificity of training effects. Ss showed training effect against Sa, but not against Pa. Conversely, Ec showed training effect against Pa, but not against Sa. We speculate that the specificity of these training effects might reflect the broad specificity of innate immune signaling pathways. However, overall, the trained immunity in *Drosophila* exhibited heterologous effects, consistent with the features of trained immunity observed in other organisms.

To further characterize the trained immunity, we focus hereafter on two experimental systems: Ml-training and Sa-challenge (MA system), and St-training and Pa-challenge (TP system).

### Persistence and removal of training bacteria

Training effects on survival rate could be due to persistence or memory of immune responses. To address this issue, we initially evaluated persistence of training bacteria after injection and measured bacteria load of Ml and St. Immediately after injection, Ml bacteria load was approximately 12,000 colony formation units (cfu) / fly in our experiments, and it gradually decreased day by day (Fig. 2A). At day 6 after injection, most of the Ml was removed from flies, and only 0.5% of the initial load (31 cfu / fly) was retained. After 12 days, Ml bacteria were not detected in fly anymore. In contrast, St bacteria load was kept at high levels and even increased to about 20 fold after 15 days (from 2,650 to 68,250 cfu / fly) (Fig. 2B). Thus, Ml and St bacteria behave differently in fly: Ml was removed, but St was persistent. A previous study [37] showed that St bacteria are incorporated into phagosomes of hemocytes and live there for a long time, and it was also suggested that St bacteria activate immune responses continuously in fly.

**Fig. 2.**
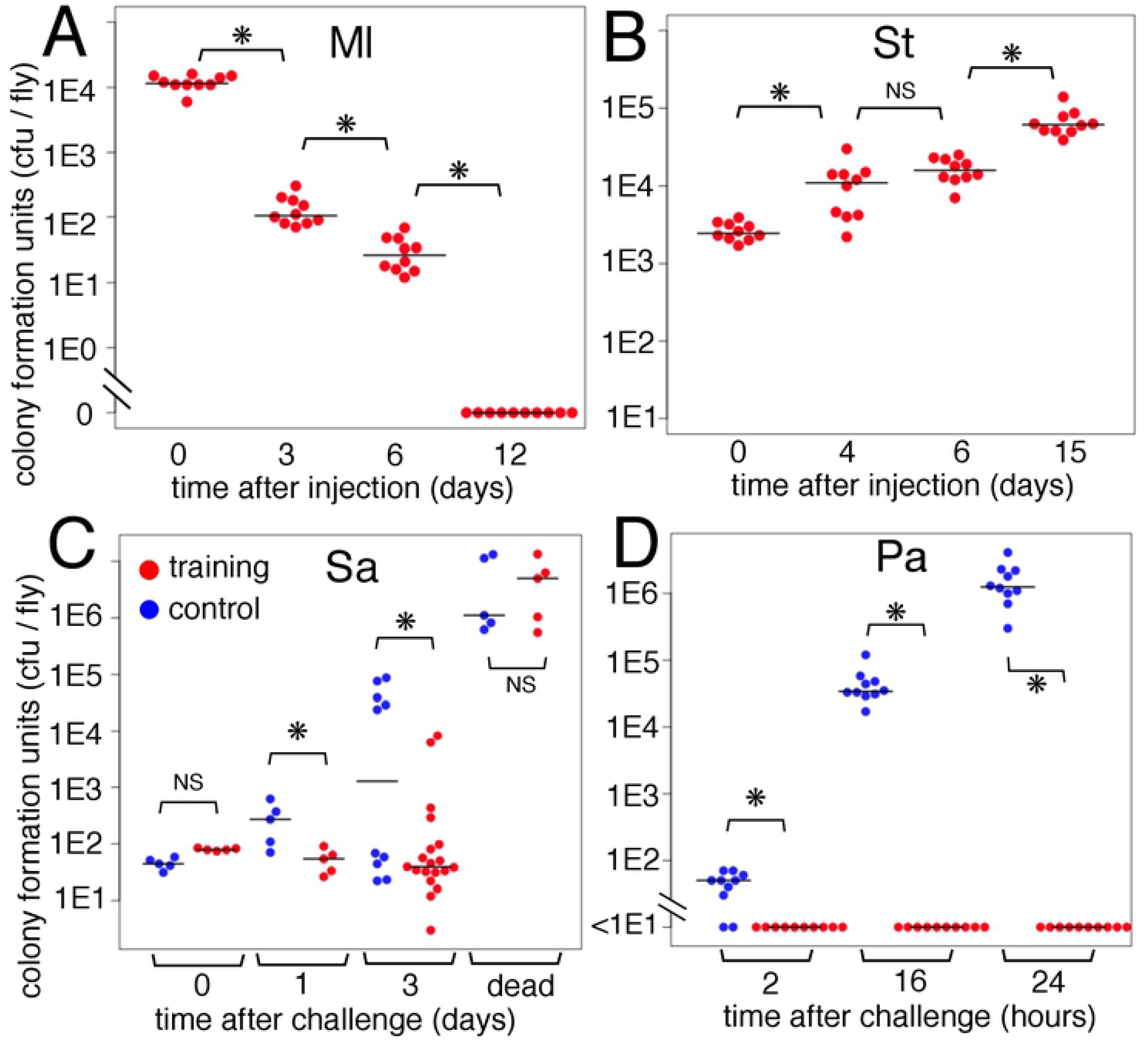
Bacteria load assay for trained immunity (A, B) Loads of training bacteria, Ml (A) and St (B). The flies were collected at given times after training injection. (C, D) Loads of challenge bacteria, Sa-GFP (C) and Pa-Kan (D). The flies were collected at the indicated times after challenge injection. “Dead” means the fly died within 24 hours before sample preparation. Asterisks and NS indicate statistically significant (p-value < 0.05) and not significant, respectively, in Kruskal-Wallis ANOVA and post-hoc Wilcoxon rank sum test.

Ml was completely removed from fly by 12 days after training. How long did the training effect of Ml lastã At day 12, the Ml-training still increased the survival of flies after Sa challenge as effectively as at day 6 (S2 Fig). This indicates that live Ml bacteria are not needed for training effects at the time of challenge. However, the training effect of Ml seemed not to be permanent, because at day 23, Ml-training did not increase fly survival after challenge.

We next asked whether immune training is detrimental for fly longevity (S3 Fig). The flies with Ml-training showed almost the same longevity as control flies. However, as shown previously [37], the flies with St-training showed significantly shorter longevity (median survival time, 32 days) than the control (59 days). Thus, persistent St bacteria is advantageous for flies by enhancing immunity during a short term but is detrimental for maintaining longevity during a long term. In contrast, we did not detect such negative effects of Ml-training.

### Clearance of challenge bacteria and survival

Life-or-death under infection would be the overall results of immunity (to remove pathogens), resistance (to protect flies from pathogen-induced damage) and other physiological states [38]. To characterize the phenomena in our survival assay, we measured the load of challenge bacteria in live and dead flies. We used antibiotic-resistant strains of challenge bacteria to distinguish them from training bacteria (see Materials and Methods). In the TP system, we observed that Pa bacteria sharply increased in the control flies, but were completely eliminated within 2 hours in the St-trained flies (Fig. 2D). This all-or-none load is consistent with the survival rate in the TP system: most of the control flies died after Pa-challenge, but most of the St-trained flies survived (Fig. 1C). Thus, St-training stimulated the immunity of flies to remove Pa.

In the MA system, Sa bacteria slowly increased in the control live flies, and at day 3 after challenge, we observed two discrete populations of the live flies that had either a high or low load of Sa bacteria (Fig. 2C). This high load was mostly close to that of the dead flies at day 3, suggesting that the fly individuals having a high load of Sa might be going to die shortly, while the flies having a low load might survive longer. After Ml-training, many flies maintained a low load of Sa, suggesting that Ml-training enhances fly immunity to remove Sa. Moreover, the Ml-trained flies at day 3 showed a relatively broad distribution of Sa load, while the dead Ml-trained flies carried a high load of Sa similar to that of the dead control flies. This result suggests that Ml-trained flies might be resistant against middle ranges of Sa load (see Discussion).

### RNA-Seq analysis of trained immunity

To address the molecular mechanism of trained immunity, we performed RNA-Seq analysis for our experimental systems. We took 3 biological replicates for each of 7 conditions in MA and TP systems: Control (Ct-Ct), Ml-training only (Ml-Ct), Sa-challenge only (Ct-Sa), Ml-training plus Sa-challenge (Ml-Sa), St-training only (St-Ct), Pa-challenge only (Ct-Pa), and St-training plus Pa-challenge (St-Pa) conditions. Total RNAs were extracted from the whole body of flies at 4 hours after challenge. Sequence reads were mapped on the *Drosophila* genome (S1 Table), and read counts of each gene were analyzed (S1 Data, S2 Data).

To obtain an overview of the transcriptome profiles of all samples, we initially performed multidimensional scaling (MDS) analysis (Fig. 3A). In the MA system, the transcriptome profile of the training only (Ml-Ct: green triangles) was similar to that of control (Ct-Ct: blue circles), while the profile of the challenge only (Ct-Sa: yellow triangles) was similar to that of the training plus challenge (Ml-Sa: red triangles). In the TP system, the transcriptome profile of the training only (St-Ct: green squares) was different to that of the control (Ct-Ct: blue circles) but was similar to that of the challenge only (Ct-Pa: yellow squares) and the training plus challenge (St-Pa: red squares). Thus, these results suggest that the transcriptome of Ml-training is close to the steady state, while the transcriptome of St-training is similar to the state of immune activation. This is consistent with the result of bacteria load: Ml was removed, while St was persistent in the flies.

**Fig. 3.**
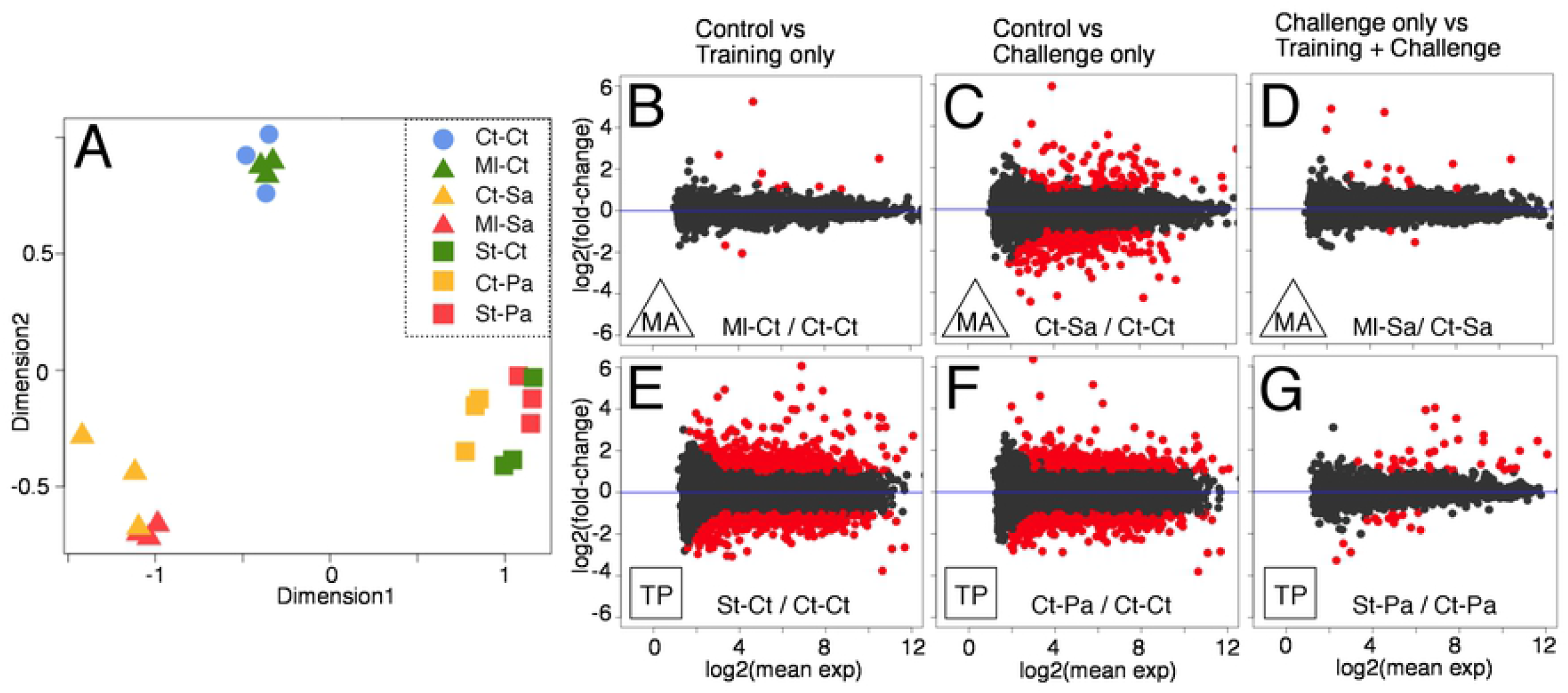
Overall features of transcriptomes (A) Multidimensional scaling analysis of transcriptomes for all samples. Distance between plots correlates to the similarity of transcriptomes between samples. (B-G) Plots of the fold-change (log2 values on Y-axis) and the mean expression (log2 values on X-axis) of the genes for pair-wise comparisons between the samples of the MA (B-D) and TP (E-G) systems. Comparisons are between control and training only (B, E), between control and challenge only (C, F), and between challenge and training plus challenge (D, G). Red dots represent DEGs showing FDR-adjusted p-value < 0.05 and the absolute value of fold-change >= 2 (see also Table 1).

Pairwise comparisons between the conditions identified differentially expressed genes (DEGs) (Table 1). In the MA system, comparisons between control (Ct-Ct) and training only (Ml-Ct) detected fewer DEGs (80 genes in total, 11 genes in fold-change >= 2) than that between control (Ct-Ct) and challenge only (Ct-Sa) (1298 genes in total, 357 genes in fold-change >= 2) (Fig. 3B, C). However, in the TP system, comparison between control (Ct-Ct) and training only (St-Ct) detected as many DEGs (5033 genes in total, 1377 genes in fold-change >= 2) as the comparison between control (Ct-Ct) and challenge only (Ct-Pa) (5037 genes in total, 1337 genes in fold-change >= 2) (Fig. 3E, F). In both systems, comparisons between challenge only (Ct-Sa or Ct-Pa) and training plus challenge (Ml-Sa or St-Pa) detected relatively small numbers of DEGs (Fig. 3D, G, Table 1). These data are consistent with the results of MDS analysis.

**Table 1.** DEGs detected from pair-wise comparisons DEGs was detected from pair-wise comparisons using a criterion (FDR-adjusted p-value < 0.05). Some DEGs were up-or down-regulated in the Y condition compared to the X condition. Some DEGs were up-or down-regulated more than 2-fold. Numbers of DEGs were indicated for each comparison.

| subject X | query Y | UP<br>Y > X | UP<br>2-fold more | DOWN<br>Y < X | DOWN<br>2-fold more | TOTAL<br>UP+DOWN |
| --- | --- | --- | --- | --- | --- | --- |
| Ct-Ct | MI-Ct | 55 | 9 | 25 | 2 | 80 |
| Ct-Ct | Ct-Sa | 516 | 98 | 782 | 259 | 1298 |
| Ct-Ct | MI-Sa | 451 | 163 | 772 | 235 | 1223 |
| MI-Ct | Ct-Sa | 637 | 87 | 860 | 306 | 1497 |
| MI-Ct | MI-Sa | 419 | 132 | 747 | 252 | 1166 |
| Ct-Sa | MI-Sa | 115 | 16 | 44 | 2 | 159 |
| Ct-Ct | St-Ct | 3006 | 887 | 2027 | 490 | 5033 |
| Ct-Ct | Ct-Pa | 3235 | 903 | 1802 | 434 | 5037 |
| Ct-Ct | St-Pa | 3134 | 942 | 2041 | 513 | 5175 |
| St-Ct | Ct-Pa | 74 | 3 | 141 | 46 | 215 |
| St-Ct | St-Pa | 25 | 0 | 54 | 17 | 79 |
| Ct-Pa | St-Pa | 113 | 35 | 50 | 14 | 163 |

### Gene expression patterns of trained immunity

To characterize patterns of gene expression in trained immunity, we calculated Z-scores (normalized deviations of expression) for DEGs (2077 genes in the MA system, 5965 genes in the TP system) and displayed the heatmap of Z-scores clustered by genes (in rows) (Fig. 4A, B). The heatmaps clearly showed the similarity of transcriptomes between Ml-training and control in the MA system, and the similarity between St-training and Pa-challenge in the TP system. These data further support the steady state for Ml-training and the immune-active state for St-training.

**Fig. 4.**
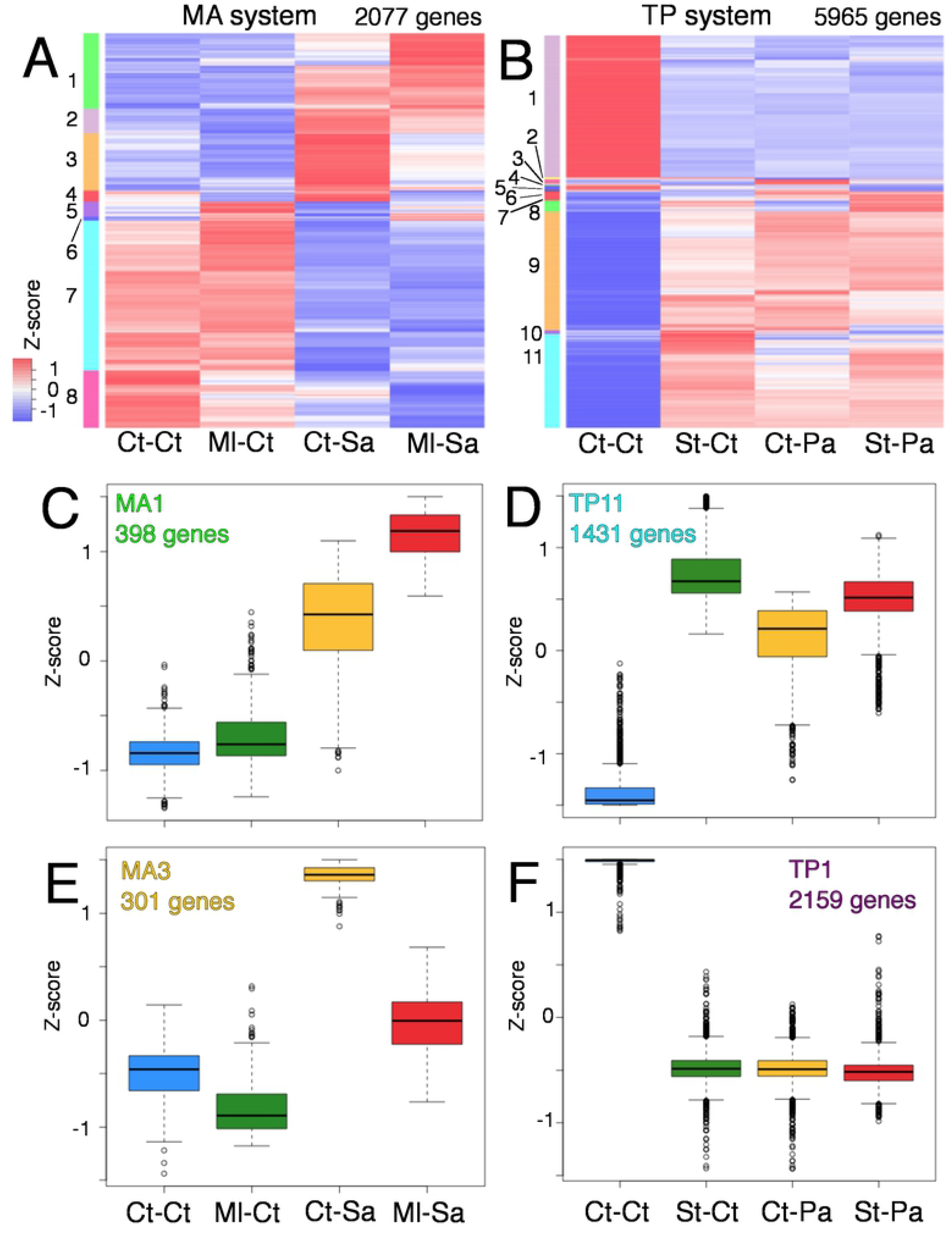
Clustering analysis of DEGs (A, B) Heat maps of z-scores for DEGs of MA (A) and TP (B) systems. Genes (rows) were ordered by clustering analysis. Numbers on the left side indicate the clustering groups. (C-F) Box plots of z-scores of DEGs categorized to MA1 (C), TP11 (D), MA3 (E), and TP1 (F) groups. The numbers of genes categorized to these groups are indicated in graphs.

Using given thresholds of clustering dendrograms, we categorized DEGs into 8 and 11 groups for the MA and TP systems, respectively (Fig. 4A, B, colored bars and numbers on the left side of heatmap), and analyzed the expression pattern of each clustering group (S3 Data, S4 Data). In the MA system, 398 genes of MA1 group were up-regulated by challenge and were further stimulated by training plus challenge (Fig. 4C). On the other hand, 301 genes of the MA3 group were up-regulated by challenge but were suppressed by training plus challenge (Fig. 4E). Thus, expression patterns of the MA1 and MA3 groups recaptured “potentiation” and “tolerance” of gene expression in trained immunity, respectively (see Introduction). In the case of the TP system, the majority of DEGs were categorized into the TP11 group (up-regulated by any stimulus) (Fig. 4D) or TP1 group (down-regulated by any stimulus) (Fig. 4F). In addition, some groups of the MA and TP systems shared similar expression patterns. For example, the genes of the MA5 and TP8 groups were up-regulated by training, but not by challenge (S4 Fig, S5 Fig). Expression patterns of other clustering groups were diverse, as shown there.

To explore biological functions of DEGs, we performed Gene Ontology (GO) analysis for each clustering group and observed significant enrichment of some GO terms (S5 Data, S6 Data). The MA1 group of the MA system was enriched for the genes related to “GO:0045087∼innate immune response” (Fig. 5). The TP11 group of the TP system was also enriched for the genes related to the same GO term. Indeed, MA1 shared 124 genes with TP11 (1.3-fold enrichment, p-value = 8.2E-4, chi-square test), and the shared genes were enriched again for the “innate immune response” (13-fold enrichment, p-value = 1.1E-10, chi-square test). Thus, a significant number of the immune-related genes displayed an expression pattern like that of the MA1 group in the MA system and the TP11 group in the TP system. Indeed, many AMP genes (e.g., *Drosomycin, Cecropin A1*, and *Attacin-A* genes) showed such expression patterns (S6 Fig), although St-training induced their expression to considerably high levels. Thus, these results suggest that Ml-training potentiates expression of the immune-related genes in response to Sa-challenge, but the Ml-training only showed the steady state of their expression. In contrast, St-training solely activates expression of these genes as Pa-challenge does. Taken together, these results demonstrate that the training effects of Ml and St would be attributed to memory and persistence of immune responses, respectively.

**Fig. 5.**
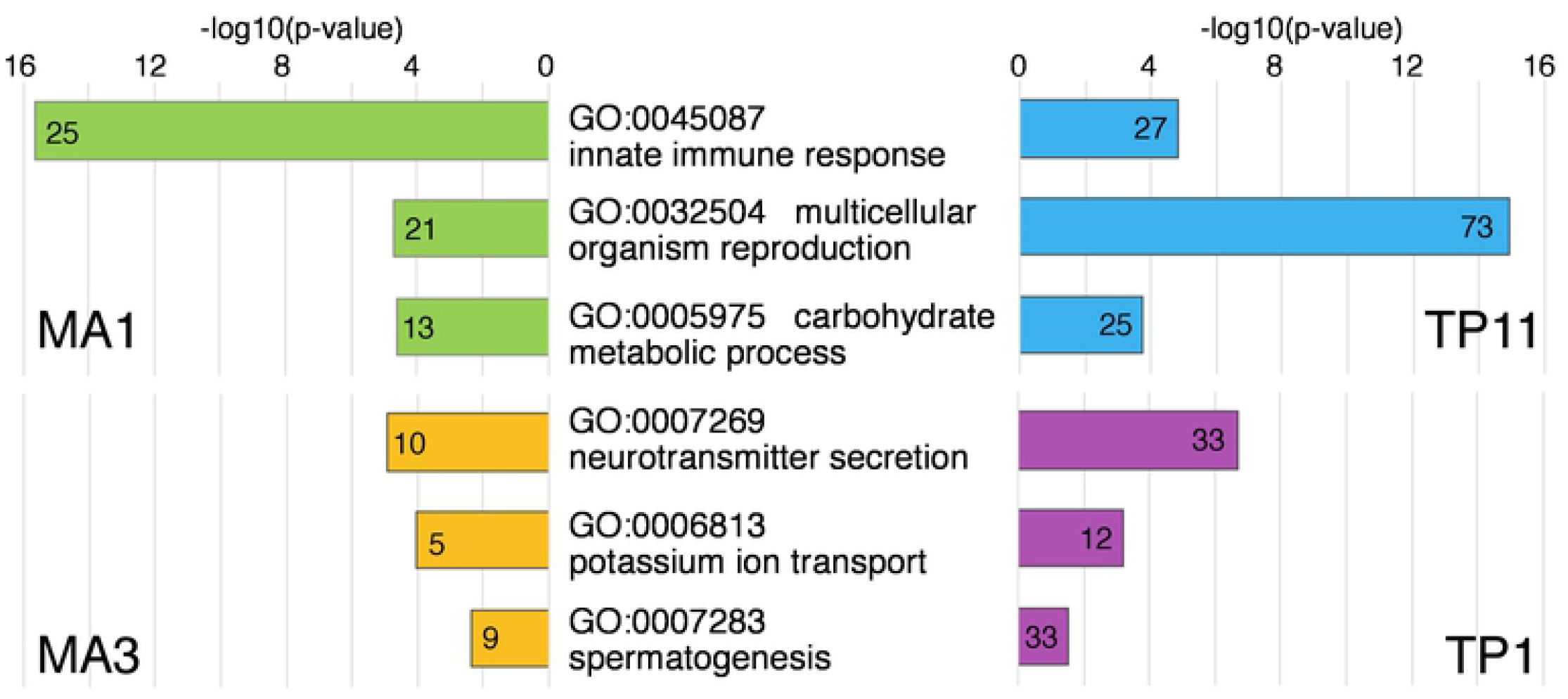
GO analysis for the clustering groups GO analysis (GOTERM_BP_DIRECT category) for each clustering group was performed (S5 Data, S6 Data). MA1 (green bars) and TP11 (blue bars) groups shared some enriched GO terms. Among them, three representative GO terms are shown here. Bar graphs show p-values of each GO term, as -log10 values. Numbers under the top of bars indicate numbers of genes matched to the GO terms. MA3 (yellow bars) and TP11 (magenta bars) groups also shared some enriched GO terms.

The MA1 and TP11 groups shared enriched GO terms for not only innate immune response, but also for multicellular organism reproduction (GO:0032504) and carbohydrate metabolic process (GO:0005975) (Fig. 5). This suggests that the genes related to innate immune response, multicellular organism reproduction, and carbohydrate metabolic process, might be regulated by comparable pathways during trained immunity. It is also possible that reproduction physiology and carbohydrate metabolism might contribute to maintaining homeostasis of fly individuals during trained immunity (see Discussion).

We found that the genes related to “GO:0045087∼innate immune response” were also enriched for other clustering groups: MA5 and TP8 (S5 Data, S6 Data). However, the expression patterns of these groups were apparently different to those of MA1 and TP11. The genes of MA5 and TP8 groups were up-regulated by training, but not by challenge (S4 Fig, S5 Fig). Therefore, the immune-related genes were controlled in several different manners during trained immunity.

MA3 and TP1 groups were enriched for genes related to “GO:0007269∼ neurotransmitter secretion” (Fig. 5). The MA3 group showed the expression patterns of immune “tolerance” (Fig. 4E), but the TP1 group was down-regulated by either training or challenge (Fig. 4F). MA3 and TP1 groups also shared the enriched GO terms: “GO:0006813∼ potassium ion transport” and “GO:0007283∼ spermatogenesis”. It is possible that these biological functions might be related to the immune tolerance in the MA system and the immune suppression in the TP system.

As other examples, the MA4, MA8, and TP1 groups were enriched for the genes related to “GO:0010906∼regulation of glucose metabolic process” (S5 Data, S6 Data). These groups share the expression pattern in which the genes are down-regulated by training regardless of challenge.

We will need further experiments to evaluate whether these genes described above function in trained immunity.

### Involvement of Ada2b in trained immunity

Our results suggest that the MA system represents immune memory, and the expression pattern of the MA1 group would recapture the potentiation of immune responses under trained immunity at the molecular level. Therefore, we focus on the mechanism governing the expression of the MA1 genes hereafter. We searched the database of publications for enrichment of the genes of the MA1 group, and found a study regarding Ada2b, a component of the chromatin modification complex [25]. This study carried out microarray analysis of *Ada2b* mutant, and identified the Ada2b-regulated genes that actually included many immune-related genes. The genes of the MA1 group were significantly enriched for the Ada2b-regulated genes (52 matches, p-value = 1.0E-44). Then, we analyzed the expression patterns of the Ada2b-dependent immune-related genes (88 genes [25]) in our RNA-Seq data and found that about half of them indeed showed the MA1-like expression pattern (step-wise activation by training and challenge) (S7 Fig). In contrast, the Ada2b-independent immune-related genes (30 genes [25]) rarely displayed such an expression pattern. As control, expression of the house-keeping genes [39] was not altered in our assay conditions. Thus, the genes enriched to the MA1 group are not merely immune-related genes, but rather the Ada2b-regulated immune-related genes. From these observations, we hypothesize that Ada2b might be involved in the trained immunity.

To evaluate our hypothesis, we examined the effects of knock down using an *Ada2b* RNAi line. Our qPCR analysis confirmed that the *Ada2b* RNAi line (da-Gal4, UAS-*Ada2b* RNAi genotype) knocked down the expression of *Ada2b* mRNA to about half of the control level (Fig. 6A). We then performed the survival assay and observed that Ml-training did not increase the survival rate of the *Ada2b* RNAi flies after Sa-challenge, while the control line (da-Gal4, UAS-GFP RNAi genotype) showed apparent training effects of Ml (Fig. 6B, C). Moreover, we observed that in the *Ada2b* mutant heterozygotes (*Ada2b* [d272] / +) [24], the effect of Ml-training was abolished (Fig. 6D, E). These results indicate that Ada2b is required for the survival enhancement in trained immunity. Further analyses are needed for clarifying the roles of Ada2b in trained immunity.

**Fig. 6.**
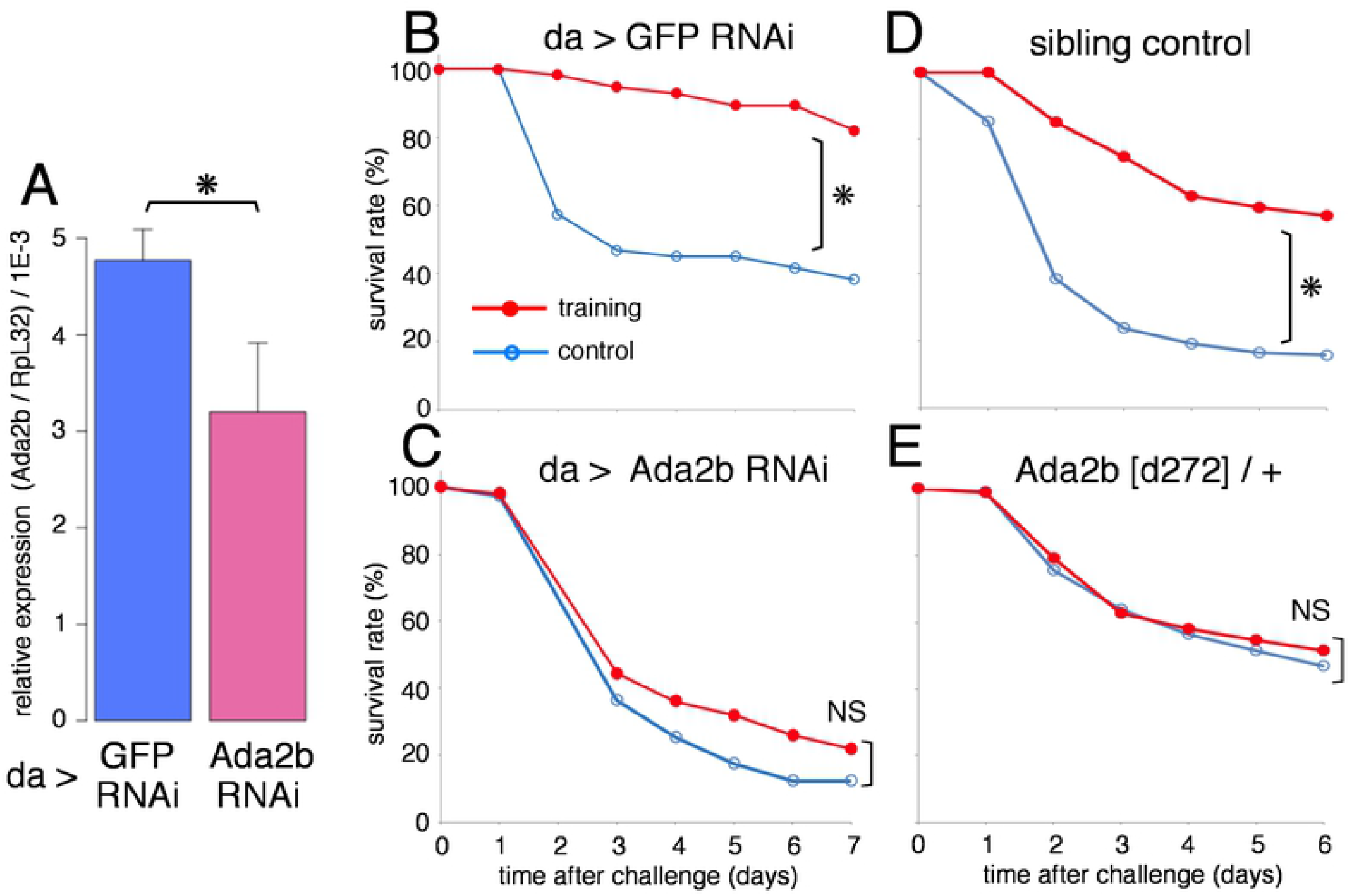
Involvement of Ada2b in trained immunity (A) qPCR analysis measured relative expression of the *Ada2b* gene (against the *RpL32* control gene). *Ada2b* RNAi significantly reduced expression of *Ada2b*. Asterisks indicate statistically significant (p-value < 0.05) in Student t-test. (B-E) Survival curves of flies under Ml-training and Sa-challenge. Red and blue lines represent the flies with and without training, respectively. (B) Control line (da > GFP RNAi), (C) *Ada2b* knock down line (da > *Ada2b* RNAi), (D) Sibling control line (w; TM6 / +), (E) *Ada2b* heterozygotes (*Ada2b* [d272] / +). Asterisks and NS indicate statistically significant (p-value < 0.05) and not significant, respectively, in Log-rank test. Numbers of flies used in these experiments are (B) 56, 57, (C) 96, 100 (D) 80, 92, (E) 97, 103 (with or without training, respectively).

## Discussion

We established the MA and TP experimental systems for detecting trained immunity in *Drosophila*. In the TP system, St bacteria persisted in flies for at least 15 days after injection, and the overall transcriptome of St-training was similar to the state of immune activation, suggesting that the training effect of St would be attributed to immune persistence. In contrast, the training effect of the MA system would be associated with immune memory, because Ml left training effects even after its removal from flies. Furthermore, the transcriptome of Ml-training was similar to that of the control, but the training enhanced the expression of the immune-related genes after Sa-challenge. Our study thus provides models of immune memory and persistence in trained immunity.

In the MA system, training effects of Ml on survival rate was dampened after 23 days. In the mammalian system, it has been shown that epigenetic information is memorized in the HSPCs, and immune memory lasts as long as the HSPCs produce peripheral myeloid cells [14,15]. In our experimental system, it is possible that hidden memory might remain at the molecular level. An another important issue in our study is which cells are responsible for trained immunity in *Drosophila*. A recent study of single-cell RNA-Seq revealed the transcriptome atlas of adult fly on the single-cell level, and identified subtypes of hemocytes and their progenitor cells [40]. This database would be valuable for identifying the cells responsible for trained immunity. Regarding the endurance of the training effect, a recent study demonstrated the transgenerational inheritance of innate immune memory in mammals [41]. Whether the trained immunity in *Drosophila* is transmitted to the next generation would be an interesting issue for future studies.

Survival of infected individuals would be determined by the combination of immunity and resistance [38]. To dissect the processes in our survival assay, we measured bacteria loads after challenge infection. In the control of the MA system, we detected two discrete fly populations having high and low loads of Sa at day 3 after challenge. This suggest that there might be a threshold of bacteria load, to which a fly resists or succumbs. Namely, when a fly succumbs to a load of bacteria, bacteria would grow rapidly to the maximum load. In the case of the Ml-training, many flies kept low loads of Sa, suggesting that Ml-training reinforces the immunity to prevent bacteria growth. Moreover, the Ml-trained flies showed a broad distribution of Sa load. This suggest that Ml-trained flies might be resistant against middle ranges of Sa load. Thus, Ml-training might potentiate both immunity and resistance of flies against Sa infection.

Transcriptome analysis revealed that the genes related to innate immunity were enriched to the MA1 and TP11 clustering groups. The expression patterns of these genes suggest that St-training induces persistence of immune activation, while Ml-training enhances immune responses after challenge via immune memory. The GO terms shared between the MA1 and TP11 groups include not only “innate immune response”, but also “multicellular organism reproduction” and “carbohydrate metabolism process”. We speculate that these genes might be regulated via comparable pathways and might be involved in trained immunity. No doubt immune responses influence many aspects of physiology, metabolism, and behaviors, and these responses altogether impact the homeostasis of the organism [20]. We thus consider it possible that reproductive physiology and carbohydrate metabolism might contribute to maintaining homeostasis under trained immunity.

In mammalian innate immune cells, it is known that immune training changes gene expression representatively via two modes. Up-regulation of gene expression under challenge conditions is enhanced or suppressed by advance training [10–13]. The genes showing potentiation and tolerance under trained immunity are enriched for the AMP genes and the pro-inflammatory genes, respectively. Thus, these studies in mammals suggest that immune training enhances the immunity to eliminate pathogens and suppresses the inflammatory response to prevent detrimental effects of immunity, and both contribute to maintaining homeostasis of the organism under trained immunity.

In the *Drosophila* MA system, the expression pattern of the MA1 group corresponds to the potentiation mode under immune training. Moreover, these genes were enriched in the innate immune response genes, including many AMP genes. Thus, the enhancement of immune-related genes under trained immunity is conserved between flies and mammals. On the other hand, the expression pattern of the MA3 group corresponds to tolerance mode under training. However, these genes were not enriched for any GO term related to immunity, but rather for “neurotransmitter secretion”. We consider it possible that the neurotransmitter might contribute to the trained immunity on the individual level, although we need further experiments to evaluate this possibility. It is known that neural networks respond to and regulate immunity [42]. For example, a recent study demonstrated that some neurons are activated by local inflammation and retrieve the inflammatory state [41]. The function of neural networks in trained immunity would be an attractive future issue.

We searched for the factor(s) involved in trained immunity of *Drosophila* and identified a candidate, Ada2b. The RNAi and mutant lines for *Ada2b* showed dampened survival enhancement via Ml-training. This result suggests that the Ada2b is involved in the trained immunity. It is known that Ada2b interacts with the HAT (histone acetyltransferase) module containing GCN5 and Ada3 [24,25,43]. The HAT module further associates with many other proteins to form large complexes (e.g., SAGA complex). Ada2b-containing complexes induce acetylation of histone H3 Lys9 (H3K9) and H3 Lys14 (H3K14) and regulate gene expression. Although we consider it possible that the Ada2b-containing complex might be involved in epigenetic regulation during trained immunity, further analysis is needed to clarify the roles of Ada2b and complexes containing it in trained immunity.

Here, we showed transcriptome features of trained immunity in *Drosophila*. However, we still do not know how gene expression is organized in spatiotemporal manners on the individual level. Moreover, we need to address the extent to which the mechanism underlying trained immunity is evolutionarily conserved. Our *Drosophila* system for trained immunity would be useful to address these issues in the future.

## Acknowledgements

We thank Drs. Chikara Kaito (Okayama Univ.), Takayuki Kuraishi (Kanazawa Univ.), Kazuhiro Iiyama (Kyushu Univ.), Naoki Hayashi (Kyoto Pharm Univ.), Nóra Zsindely (Univ. of Szeged), and Mattias Mannervik (Stockholm Univ.) for providing bacteria strains and fly strains to us. We thank the Bloomington Drosophila Stock Center, Vienna Drosophila Resource Center, Kyoto Stock Center, and NIG Stock Center for providing various fly stocks. We also thank the members of the Kurata Laboratory for discussions and suggestions. This work was supported by JSPS KAKENHI Grant Number 16H06279 (PAGS), 17K07239 and 19H03365. Computations were partially performed on the NIG supercomputer at ROIS National Institute of Genetics.

## References

1. Netea MG, Domínguez-Andrés J, Barreiro LB, Chavakis T, Divangahi M, et al. (2020) Defining trained immunity and its role in health and disease. Nat Rev Immunol 20: 375–388. Available: https://www.nature.com/articles/s41577-020-0285-6. Accessed 25 August 2021.

2. Netea MG, Schlitzer A, Placek K, Joosten LAB, Schultze JL (2019) Innate and Adaptive Immune Memory: an Evolutionary Continuum in the Host’s Response to Pathogens. Cell Host Microbe 25: 13–26. Available: https://linkinghub.elsevier.com/retrieve/pii/S1931312818306334.

3. Kaufmann E, Sanz J, Dunn JL, Khan N, Mendonça LE, et al. (2018) BCG Educates Hematopoietic Stem Cells to Generate Protective Innate Immunity against Tuberculosis. Cell 172: 176–190.e19. Available: http://linkinghub.elsevier.com/retrieve/pii/S0092867417315118. Accessed 12 January 2018.

4. Cirovic B, de Bree LCJ, Groh L, Blok BA, Chan J, et al. (2020) BCG Vaccination in Humans Elicits Trained Immunity via the Hematopoietic Progenitor Compartment. Cell Host Microbe 28: 322–334. Available: https://doi.org/10.1016/j.chom.2020.05.014. Accessed 17 August 2020.

5. Muñoz N, Van Maele L, Marqués JM, Rial A, Sirard J-C, et al. (2010) Mucosal Administration of Flagellin Protects Mice from Streptococcus pneumoniae Lung Infection. Infect Immun 78: 4226–4233. Available: https://journals.asm.org/doi/10.1128/IAI.00224-10.

6. Milutinović B, Kurtz J (2016) Immune memory in invertebrates. Semin Immunol 28: 328–342. Available: http://www.sciencedirect.com/science/article/pii/S1044532316300434. Accessed 19 January 2018.

7. Kachroo A, Robin GP (2013) Systemic signaling during plant defense. Curr Opin Plant Biol 16: 527–533. doi:10.1016/J.PBI.2013.06.019.

8. Gourbal B, Pinaud S, Beckers GJM, Van Der Meer JWM, Conrath U, et al. (2018) Innate immune memory: An evolutionary perspective. Immunol Rev 283: 21–40. Available: http://doi.wiley.com/10.1111/imr.12647.

9. Divangahi M, Aaby P, Khader SA, Barreiro LB, Bekkering S, et al. (2021) Trained immunity, tolerance, priming and differentiation: distinct immunological processes. Nat Immunol 22: 2–6. Available: http://www.nature.com/articles/s41590-020-00845-6.

10. Quintin J, Saeed S, Martens JHA, Giamarellos-Bourboulis EJ, Ifrim DC, et al. (2012) Candida albicans Infection Affords Protection against Reinfection via Functional Reprogramming of Monocytes. 223–232 p. Available: http://www.sciencedirect.com/science/article/pii/S1931312812002326. Accessed 2 August 2017.

11. Saeed S, Quintin J, Kerstens HHD, Rao NA, Aghajanirefah A, et al. (2014) Epigenetic programming of monocyte-to-macrophage differentiation and trained innate immunity. Science (80-) 345. Available: http://science.sciencemag.org/content/345/6204/1251086. Accessed 6 April 2017.

12. Foster SL, Hargreaves DC, Medzhitov R (2007) Gene-specific control of inflammation by TLR-induced chromatin modifications. Nature 447: 972–978. Available: http://dx.doi.org/10.1038/nature05836. Accessed 16 May 2016.

13. Zhang Q, Cao X (2019) Epigenetic regulation of the innate immune response to infection. Nat Rev Immunol 19: 417–432. Available: http://www.nature.com/articles/s41577-019-0151-6.

14. de Laval B, Maurizio J, Kandalla PK, Brisou G, Simonnet L, et al. (2020) C/EBPβ-Dependent Epigenetic Memory Induces Trained Immunity in Hematopoietic Stem Cells. Cell Stem Cell. Available: https://linkinghub.elsevier.com/retrieve/pii/S1934590920300175.

15. Mitroulis I, Ruppova K, Wang B, Chen L-S, Grzybek M, et al. (2018) Modulation of Myelopoiesis Progenitors Is an Integral Component of Trained Immunity. Cell 172: 147–161.e12. Available: http://linkinghub.elsevier.com/retrieve/pii/S0092867417313855. Accessed 12 January 2018.

16. Christ A, Günther P, Lauterbach MAR, Duewell P, Biswas D, et al. (2018) Western Diet Triggers NLRP3-Dependent Innate Immune Reprogramming. Cell 172: 162–175.e14. Available: http://linkinghub.elsevier.com/retrieve/pii/S0092867417314939. Accessed 12 January 2018.

17. Bekkering S, Arts RJW, Novakovic B, Kourtzelis I, van der Heijden CDCC, et al. (2018) Metabolic Induction of Trained Immunity through the Mevalonate Pathway. Cell 172: 135–146.e9. Available: http://linkinghub.elsevier.com/retrieve/pii/S0092867417313727. Accessed 12 January 2018.

18. Lemaitre B, Hoffmann J (2007) The Host Defense of Drosophila melanogaster. Annu Rev Immunol 25: 697–743. Available: http://www.annualreviews.org/doi/10.1146/annurev.immunol.25.022106.141615.

19. Kimbrell DA, Beutler B (2001) The evolution and genetics of innate immunity. Nat Rev Genet 2: 256–267. Available: https://www.nature.com/articles/35066006. Accessed 15 November 2021.

20. Buchon N, Silverman N, Cherry S (2014) Immunity in Drosophila melanogaster — from microbial recognition to whole-organism physiology. Nat Rev Immunol 14: 796–810. Available: http://dx.doi.org/10.1038/nri3763. Accessed 25 November 2014.

21. Boman HG, Nilsson I, Rasmuson B (1972) Inducible Antibacterial Defence System in Drosophila. Nature 237: 232–235. Available: http://www.nature.com/doifinder/10.1038/237232a0.

22. Pham LN, Dionne MS, Shirasu-Hiza M, Schneider DS (2007) A specific primed immune response in Drosophila is dependent on phagocytes. PLoS Pathog 3: e26. Available: http://journals.plos.org/plospathogens/articleãid=10.1371/journal.ppat.0030026. Accessed 13 April 2016.

23. Christofi T, Apidianakis Y, Christofi T, Apidianakis Y (2013) Drosophila immune priming against Pseudomonas aeruginosa is short-lasting and depends on cellular and humoral immunity. F1000Research 2. Available: http://f1000research.com/articles/2-76/v1. Accessed 14 July 2016.

24. Pankotai T, Komonyi O, Bodai L, Újfaludi Z, Muratoglu S, et al. (2005) The Homologous Drosophila Transcriptional Adaptors ADA2a and ADA2b Are both Required for Normal Development but Have Different Functions. Mol Cell Biol 25: 8215–8227. Available: http://mcb.asm.org/. Accessed 20 May 2021.

25. Zsindely N, Pankotai T, Ujfaludi Z, Lakatos D, Komonyi O, et al. (2009) The loss of histone H3 lysine 9 acetylation due to dSAGA-specific dAda2b mutation influences the expression of only a small subset of genes. Nucleic Acids Res 37: 6665–6680. Available: https://academic.oup.com/nar/article-lookup/doi/10.1093/nar/gkp722.

26. Basset A, Khush RS, Braun A, Gardan L, Boccard F, et al. (2000) The phytopathogenic bacteria Erwinia carotovora infects Drosophila and activates an immune response. Proc Natl Acad Sci 97: 3376–3381. Available: http://www.pnas.org/cgi/doi/10.1073/pnas.97.7.3376.

27. Sastalla I, Chim K, Cheung GYC, Pomerantsev AP, Leppla SH (2009) Codon-Optimized Fluorescent Proteins Designed for Expression in Low-GC Gram-Positive Bacteria. Appl Environ Microbiol 75: 2099–2110. Available: https://journals.asm.org/doi/10.1128/AEM.02066-08.

28. Chieda Y, Iiyama K, Lee JM, Kusakabe T, Yasunaga-Aoki C, et al. (2011) Virulence of an exotoxin A-deficient strain of Pseudomonas aeruginosa toward the silkworm, Bombyx mori. Microb Pathog 51: 407–414. Available: http://linkinghub.elsevier.com/retrieve/pii/S088240101100163X. Accessed 30 August 2017.

29. Francis KP, Yu J, Bellinger-Kawahara C, Joh D, Hawkinson MJ, et al. (2001) Visualizing Pneumococcal Infections in the Lungs of Live Mice Using Bioluminescent Streptococcus pneumoniaeTransformed with a Novel Gram-Positive luxTransposon. Infect Immun 69: 3350–3358. Available: https://iai.asm.org/content/69/5/3350.

30. Apidianakis Y, Rahme LG (2009) Drosophila melanogaster as a model host for studying Pseudomonas aeruginosa infection. Nat Protoc 4: 1285–1294. Available: http://www.nature.com/doifinder/10.1038/nprot.2009.124. Accessed 22 June 2016.

31. Martin M (2011) Cutadapt removes adapter sequences from high-throughput sequencing reads. EMBnet.journal 17: 10. Available: http://journal.embnet.org/index.php/embnetjournal/article/view/200.

32. Kim D, Langmead B, Salzberg SL (2015) HISAT: a fast spliced aligner with low memory requirements. Nat Methods 2015 124 12: 357–360. Available: https://www.nature.com/articles/nmeth.3317. Accessed 8 December 2021.

33. Anders S, Pyl PT, Huber W (2015) HTSeq--a Python framework to work with high-throughput sequencing data. Bioinformatics 31: 166–169. Available: https://academic.oup.com/bioinformatics/article-lookup/doi/10.1093/bioinformatics/btu638.

34. Huang DW, Sherman BT, Lempicki RA (2009) Systematic and integrative analysis of large gene lists using DAVID bioinformatics resources. Nat Protoc 4: 44–57. Available: http://www.ncbi.nlm.nih.gov/pubmed/19131956. Accessed 9 July 2014.

35. Lyne R, Smith R, Rutherford K, Wakeling M, Varley A, et al. (2007) FlyMine: An integrated database for Drosophila and Anopheles genomics. Genome Biol 8: 1–16. Available: https://genomebiology.biomedcentral.com/articles/10.1186/gb-2007-8-7-r129. Accessed 8 December 2021.

36. Larkin A, Marygold SJ, Antonazzo G, Attrill H, dos Santos G, et al. (2021) FlyBase: updates to the Drosophila melanogaster knowledge base. Nucleic Acids Res 49: D899–D907. Available: https://academic.oup.com/nar/article/49/D1/D899/5997437. Accessed 8 December 2021.

37. Shinzawa N, Nelson B, Aonuma H, Okado K, Fukumoto S, et al. (2009) p38 MAPK-Dependent Phagocytic Encapsulation Confers Infection Tolerance in Drosophila. Cell Host Microbe 6: 244–252. Available: http://linkinghub.elsevier.com/retrieve/pii/S1931312809002583. Accessed 1 September 2017.

38. Louie A, Song KH, Hotson A, Thomas Tate A, Schneider DS (2016) How Many Parameters Does It Take to Describe Disease Tolerance? PLOS Biol 14: e1002435. Available: http://journals.plos.org/plosbiology/articleãid=10.1371/journal.pbio.1002435#pbi o.1002435.s003. Accessed 29 April 2016.

39. De Ferrari L, Aitken S (2006) Mining housekeeping genes with a Naive Bayes classifier. BMC Genomics 7: 277. Available: https://bmcgenomics.biomedcentral.com/articles/10.1186/1471-2164-7-277. Accessed 27 April 2021.

40. Li H, Janssens J, Waegeneer M De, Kolluru SS, Davie K, et al. (2021) Fly Cell Atlas: a single-cell transcriptomic atlas of the adult fruit fly. bioRxiv: 2021.07.04.451050. Available: https://www.biorxiv.org/content/10.1101/2021.07.04.451050v1. Accessed 3 December 2021.

41. Koren T, Yifa R, Amer M, Krot M, Boshnak N, et al. (2021) Insular cortex neurons encode and retrieve specific immune responses. Cell. Available: http://www.ncbi.nlm.nih.gov/pubmed/34752731. Accessed 16 November 2021.

42. Chu C, Artis D, Chiu IM (2020) Neuro-immune Interactions in the Tissues. Immunity 52: 464–474. doi:10.1016/J.IMMUNI.2020.02.017.

43. Torres-Zelada EF, Weake VM (2021) The Gcn5 complexes in Drosophila as a model for metazoa. Biochim Biophys Acta - Gene Regul Mech 1864: 194610. doi:10.1016/J.BBAGRM.2020.194610.

